# Simultaneous Targeting of XPO1 and BCL2 as an Effective Treatment Strategy for Double-Hit Lymphoma

**DOI:** 10.1101/688093

**Authors:** Yuanhui Liu, Nancy G. Azizian, Yaling Dou, Lan V. Pham, Yulin Li

## Abstract

Double-hit lymphoma (DHL) is among the most aggressive and chemoresistant lymphoma subtypes. DHLs carry genomic abnormalities in MYC, BCL2 and/or BCL6 oncogenes. Due to simultaneous overexpression of these driver oncogenes, DHLs are highly resistant to the frontline therapies. Most DHLs overexpress both MYC and BCL2 driver oncogenes concurrently. We reasoned that simultaneous suppression of the two driver oncogenes would be more effective in eradicating DHLs than inactivation of single oncogene. XPO1 is a receptor for nuclear cytoplasmic transport of protein and RNA species. Recently, XPO1 inhibition was shown to downregulate MYC expression in several cancer cell lines. We therefore examined the role of XPO1 as a therapeutic target in suppressing MYC function and the potential synergistic effects of simultaneous suppression of XPO1 and BCL2 in the treatment of DHL. Here, we demonstrate that XPO1 inhibition abrogates MYC protein expression and induces massive tumor cell apoptosis. Combined use of XPO1 and BCL2 inhibitors is highly effective in eradicating DHL cells in cell culture. Notably, in a mouse model of DHL bearing primary tumor cells derived from lymphoma patients, combined treatment with XPO1 and BCL2 inhibitors blocks tumor progression, prevents brain metastasis, and extends host survival. Thus, our study confirms the simultaneous targeting of MYC and BCL2 driver oncogenes through the combined use of XPO1 and BCL2 inhibitors as a unique approach for the treatment of DHLs.

## Introduction

Double-hit lymphoma (DHL) is a subtype of Non-Hodgkin’s Lymphoma (NHL) with genomic abnormalities in MYC and BCL2 (and less frequently BCL6), leading to the overexpression of these driver oncogenes. The prognosis for the majority of NHLs has improved significantly in the past decades due to the development of chemotherapy, targeted therapy, and immunotherapy. In comparison, DHL remains highly resistant and refractory to the first line immunochemotherapy, R-CHOP, with a 5-year overall survival rate of 30% [1, 2].

The co-existence of multiple oncogenic events in DHLs, including MYC, BCL2, and BCL6, provides an opportunity for combined targeted therapy. As the two common drivers of DHL, MYC and BCL2 cooperate in lymphomagenesis and tumor maintenance. A combined therapy aimed at targeting both MYC and BCL2 may arguably be more effective than suppressing either MYC or BCL2 alone in eradicating the tumors cells [3, 4]. Among several BCL2 inhibitors, ABT199, was developed and tested in clinical trials, and approved by FDA for the treatment of chronic lymphocytic leukemia [5]. In contrast, direct targeting of MYC has proven challenging due to its structural property as a transcription factor.

XPO1 is an adaptor of nuclear export for many protein and RNA species. Recent studies suggest that XPO1 may regulate nuclear export of mRNAs encoding several oncoproteins, such as MYC, BCL2, Cyclin D1, and PIM1 [6–8]. Moreover, suppression of XPO1 with selective inhibitors of nuclear export (SINEs) has been shown to downregulate MYC expression in several tumor types [9–12]. We therefore hypothesize that tumor cells with concurrent MYC and BCL2 overexpression, such as DHL, may be effectively targeted through the combined treatment with XPO1 and BCL2 inhibitors.

Here, we demonstrate that XPO1 inhibition in DHL tumor cells abrogates MYC protein expression. Furthermore, combined suppression of XPO1 and BCL2 synergizes to induce massive cell death in DHL tumor cells *in vitro*. Most importantly, combination therapy with XPO1 and BCL2 inhibitors blocks tumor progression and metastasis, and extends survival in a mouse model bearing patient-derived DHL tumors. Our data suggests that combined targeting of XPO1/BCL2 can be a robust therapeutic strategy for DHLs carrying both MYC and BCL2 driver oncogenes.

## Materials and Methods

### Cell lines and chemical reagents

DHL cell lines RC, SU-DHL-4 (DHL4), SU-DHL-6 (DHL6), SU-DHL-10 (DHL10) and Toledo were purchased from the ATCC. DB, SU-DHL-5 (DHL5), OCI-LY19, Will2, WSU, Val and U2932 were gifts from Dr. Lan Pham at the University of Texas MD Anderson Cancer Center. Cells were cultured in RPMI 1640, supplemented with 20% FBS (except for RC, DHL4, and Toledo, supplemented with 10% FBS), and 1% penicillin/streptomycin in a 5% CO2 humidified incubator. ABT199 (Venetoclax) was obtained from Houston Methodist Pharmacy. KPT8602 and KPT330 were purchased from Selleckchem. Carfilzomib was purchased from Cayman Chemical.

### IC50 determination

Cells were plated at 2×10^4^ cells in 100 μl culture medium per well in 96-well plates. Drugs were added the next day in 4 replicates. Following five days of drug treatment (except for DHL10, which was treated for three days), cell viability was assessed using the CellTiter-Glo^®^ 2.0 Cell Viability Assay (Promega) according to the manufacturer’s instructions. Data analyses and calculation of median effective dose (IC50) were carried out using GraphPad Prism 8 (GraphPad).

### Western blot analysis

DHL cells were lysed on ice in CelLytic™ MT Cell Lysis Reagent (Sigma) supplemented with protease and phosphatase inhibitors. Protein concentration was determined with Pierce™ BCA Protein Assay Kit (Thermo Fisher). Protein lysates were resolved on SDS-PAGE gels and transferred to PVDF membranes. The following antibodies were used for the Western blot analysis: MYC, cleaved PARP, Caspase 3, BCL2, XPO1, MCL1, BIM, BCL-XL, Lamin B1, and GAPDH (Cell Signaling); MYC (Abcam); β-Tubulin and β-Actin (Proteintech).

### Cellular Fractionation and quantitative real-time PCR

Cytoplasmic and nuclear RNA were isolated and purified using the RNA Subcellular Isolation Kit (Active Motif) following the manufacturer’s protocol. RNA binding and purification was performed using RNeasy Mini Kit (Qiagen). Total RNA was extracted with RNeasy Mini Kit (Qiagen). cDNA was synthesized using the Verso cDNA Synthesis Kit (Thermo Scientific) according to the manufacturer’s instructions. qPCR analysis was performed on 7500 Real Time PCR System using SYBR Green (Applied Biosystems). Transcript levels were normalized to GAPDH, and relative gene expression was determined with ddCt method. Primer sequences are available upon request.

### Flow cytometry

The PE Annexin-V Apoptosis Detection Kit (BD Biosciences) was used to detect apoptosis following manufacturer’s instruction. Briefly, cells were suspended in 150 μl of binding buffer and mixed with 5 μl of FITC-conjugated Annexin-V and 7-AAD, followed by incubation at room temperature for 15 min in the dark. The stained cells were analyzed using a BD FACS Fortessa flow cytometer (BD Bioscience). All flow cytometric data were analyzed with Flowjo software (Tree Star).

### Immunohistochemistry

Mice were treated for five days. Subsequently, spleen tumors were collected, formalin-fixed, and paraffin-embedded. 4 micrometer thick sections were subjected to H&E and immunohistochemistry staining. For IHC, MYC (Abcam, ab32072), Ki67 (Abcam, ab16667), and Cleaved caspase 3 (CST, 9661) antibodies were used. Pictures were taken with 40x objectives on a Leica DMi8 microscope.

### In vivo therapeutic studies

Animal studies and experimental protocols were approved by Institutional Animal Care and Use Committee at Houston Methodist Research Institute (IACUC approval number AUP-1117-0053). All experimental methods were performed in accordance with the relevant national and institutional guidelines and regulations. The DHL PDX sample (DFBL-69487-V3-mCLP) was obtained from the Public Repository of Xenografts (PRoXe), which collects PDX samples from patients with informed consent. Cells (10^6^) were engrafted via tail vein injection into 6- to 8-week-old NSG mice. The animals were monitored once a week by whole-body imaging on an IVIS Lumina III platform. Treatment started upon the appearance of measurable tumors. Tumor volumes were assessed from the start of the treatment 2 times per week using IVIS imaging system. The following treatment schemes were used: daily oral gavage with ABT199 (50mg/kg) and KPT8602 (7.5mg/kg) for 5 successive days, followed by 2 days off for three weeks. ABT199 tablets were dissolved in water by sonication; KPT8602 was dissolved in 0.5% methylcellulose plus 1% Tween-80 as reported [13–15]. BLI data were analyzed with the Living Image software, version 4.2 (Caliper Life Sciences). Animal studies were carried out in accordance with IACUC approval at Houston Methodist Research Institute (AUP-1117-0053).

### Statistical analysis

Two-tailed Student’s t test was used to analyze the quantitative PCR data for mRNA expression. Cell death rates among different treatment groups were analyzed using ANOVA with a Tukey’s test. Results were presented as mean□±□standard deviation. Animal survival in different groups was compared by Kaplan-Meier analysis with Log-rank (Mantel-Cox) test.

## Results

### XPO inhibition abrogates MYC protein expression and induces apoptosis in DHLs

First, we examined whether XPO1 inhibition affects MYC protein levels in DHL cell lines. Treatment with the XPO1 inhibitor, KPT8602, led to a significant decrease in MYC protein expression in the majority of DHLs in our diverse cell line panel (**Fig 1A**). Three DHL cell lines (SU-DHL4, Toledo, and SU-DHL6) were selected to further examine MYC regulation by XPO1 inhibitors [16]. Treatment with two specific XPO1 inhibitors, KPT330 and KPT8602, points to a dose- and time-dependent downregulation of MYC protein expression in all three cell lines (**Fig 1B-C**, and **Supplementary Fig S1**). These results establish that XPO1 inhibition effectively abrogates MYC protein expression in DHL tumor cells. Decreased MYC protein level was followed by changes in gene expression of MYC downstream targets (**Fig 1D-E**). Genes known to be upregulated by MYC, including ENO1, APEX1, RPL3, RPS5, SRM and nucleolin [17], were significantly downregulated upon XPO1 inhibition. In contrast, genes reportedly suppressed by MYC, such as HBP1, P27, and P15, were upregulated upon treatment with XPO1 inhibitors. The abrogation of MYC protein expression by XPO1 inhibition was accompanied by induction of apoptosis, as manifested by the cleavage of PARP and Caspase-3 (**Supplementary Fig S1**). We concluded that XPO1 suppression abrogates the function of MYC oncogene and induces mass apoptosis in DHL tumor cells.

**Figure 1.**
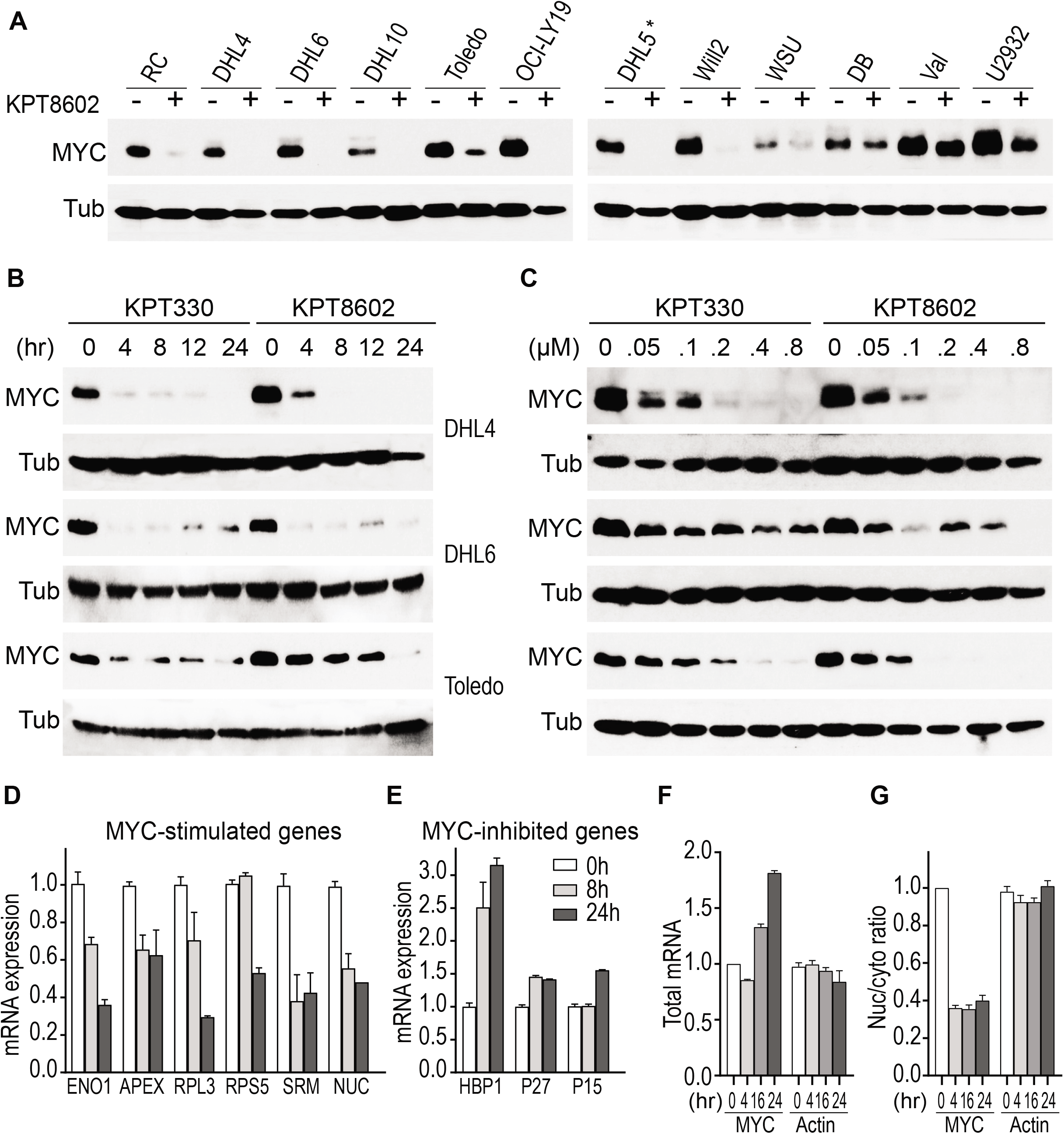
XPO1 inhibitors suppress MYC protein expression in DHLs. (A) Changes in MYC protein level upon treatment with KPT8602 (1 μM for 24 h). Asterisk (*) indicates that DHL5 is not considered a DHL line. (B) Time-dependent modulation of MYC protein in three DHL cell lines upon exposure to KPT330 or KPT8602 for 4, 8, 12 and 24 h. Cells were treated with drug concentration of 0.5 μM for DHL4/Toledo and 1 μM for DHL6. (C) Dose-dependent reduction of MYC protein level upon XPO1 inhibition. Cells were treated with 0 to 0.8 μM KPT330 or KPT8602 for 24 h. β-tubulin (Tub) serves as a loading control. (D-E) Changes in transcript levels of MYC targets upon XPO1 inhibition. DHL6 was treated with KPT8602 at 1 μM for 8 and 24 hours. (F) Total mRNA and (G) nuclear to cytoplasmic ratios of MYC and β-tubulin transcripts in DHL6 treated with 1 μM of KPT8602 for 4, 16 or 24 h. All mRNA values were normalized to GAPDH.

XPO1 has been reported to mediate nuclear export of several mRNAs encoding oncoproteins, including MYC, BCL2, and PIM1. Therefore, XPO1 inhibition may potentially reduce MYC protein expression through the nuclear retention of MYC mRNA. To test whether XPO1 inhibition affects nuclear export of MYC mRNA, we examined the distribution of MYC transcripts by cellular fractionation followed by quantitative realtime PCR. Compartmental separation of the nuclear versus cytoplasmic fractions was well achieved as shown by quantification of GAPDH and laminin B1 proteins, and Neat1 transcript in each compartment [18] (**Supplementary Fig S2C-D**). Interestingly, MYC nuclear to cytoplasmic ratio decreased by 50% upon XPO1 inhibition, while its mRNA abundance in the whole cell extract increased by two folds (**Fig 1F-G**). The nuclear to cytoplasmic ratio of BCL2 but BCL6 increased modestly at 16 hours of posttreatment. The level and nuclear/cytoplasmic ratio of beta-actin, used as an internal control, was not affected by XPO1 inhibition (**Supplementary Fig S2A-D**). This observation indicates that XPO1 inhibition may not result in a preferential nuclear retention of MYC mRNA. Therefore, the drastic downregulation of MYC protein expression upon XPO1 inhibition cannot be explained by changes in the levels and/or transport of MYC mRNA. We further ruled out the role of protein degradation as a contributing factor, as treatment with a proteasome inhibitor, carfilzomib, did not affect MYC shutdown by XPO1 inhibition (**Supplementary Fig S2E**). Taken together, these data strongly suggest that the effective downregulation of MYC protein by XPO1 inhibition may occur at the level of protein translation.

### XPO1 inhibition synergizes with BCL2 inhibition in vitro in killing DHL tumor cells

The compelling downregulation of MYC protein expression by XPO1 inhibition suggests the possibility to target MYC and BCL2 driver oncogenes in DHL by combining XPO1 and BCL2 inhibitors. To test this notion, we first determined the IC50 values for two targeted agents, ABT199 (a BCL2 inhibitor), and KPT8602 (an XPO1 inhibitor) in a panel of DHL cell lines. Among the DHL cell lines examined, some were resistant to ABT199 treatment, while others were moderately resistant to KPT8602. However, to our surprise, none of the DHL cell lines were resistant to both, lending experimental support to the combinatorial use of XPO1 and BCL2 inhibitors to eradicate DHL tumor cells (**Fig 2A-C, Supplementary Fig S3A-B**). Further *in vitro* examination of the drug combination using additional DHL cell lines demonstrated a strong synergy in cell death induction (**Fig 3A, Supplementary Fig S3C-D**), while treatment with single agent showed moderate effects. The apoptotic event resulting from the combined treatment was accompanied by enhanced PARP and Caspase 3 cleavage compared to treatment with either single agents (**Fig 3B-D**). The apoptosis phenotype induced by synergistic drug combination was further confirmed by flow cytometric analysis following 7AAD/Annexin V staining (**Fig 3E**). MCL1 is known to play an important role in regulating the sensitivity of tumor cells to BCL2 inhibitors as MCL1 overexpression is associated with resistance to ABT199 [19, 20]. Next, we examined whether XPO1 inhibition enhances the sensitivity of tumors cells to ABT199 by downregulating MCL1. Indeed, MCL1 downregulation was observed in 8 out of 12 DHL cell lines treated with XPO1 inhibitor KPT8602 (**Supplementary Fig S1A**). In DHL6 cells, the drug combination did not decrease MCL1 protein level but induced BIM protein expression (**Supplementary Fig S4A-C**). These observations may explain the synergistic effect between XPO1 and BCL2 inhibitors in inducing apoptosis of DHL tumor cells.

**Figure 2.**
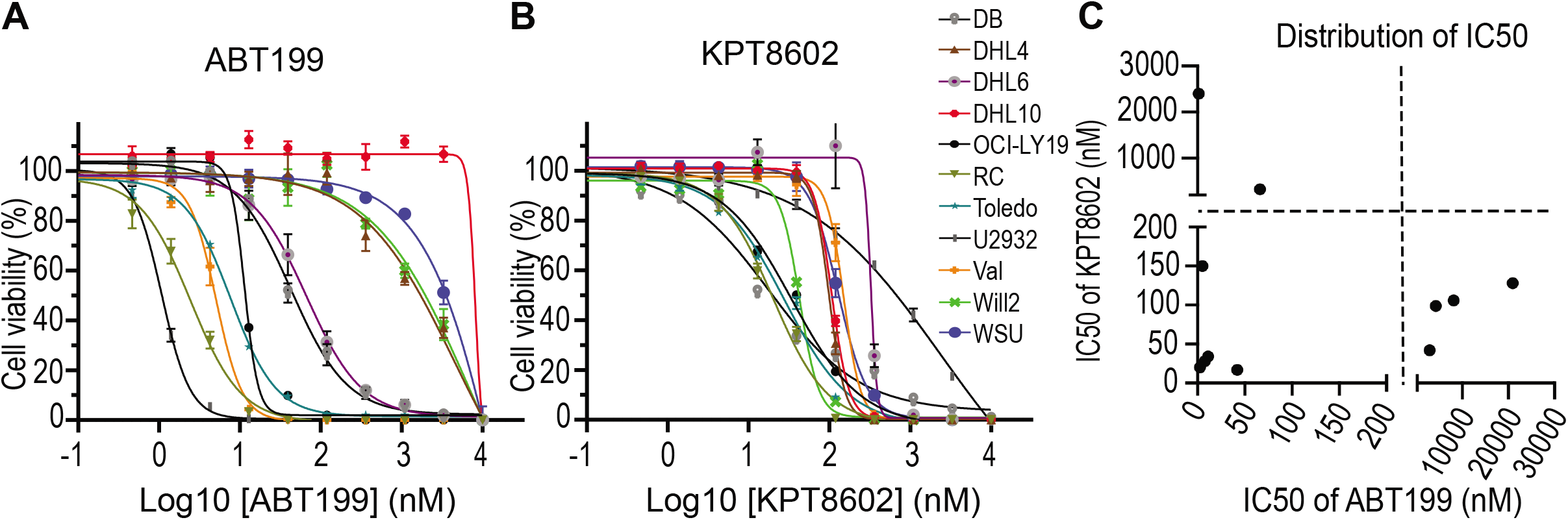
Determination of IC50 values for ABT199 and KPT8602 in a panel of DHL cell lines. (A) IC50 values for ABT199. (B) IC50 values for KPT8602. (C) IC50 values for ABT199 and KPT8602 in a panel of DHL cell lines.

**Figure 3.**
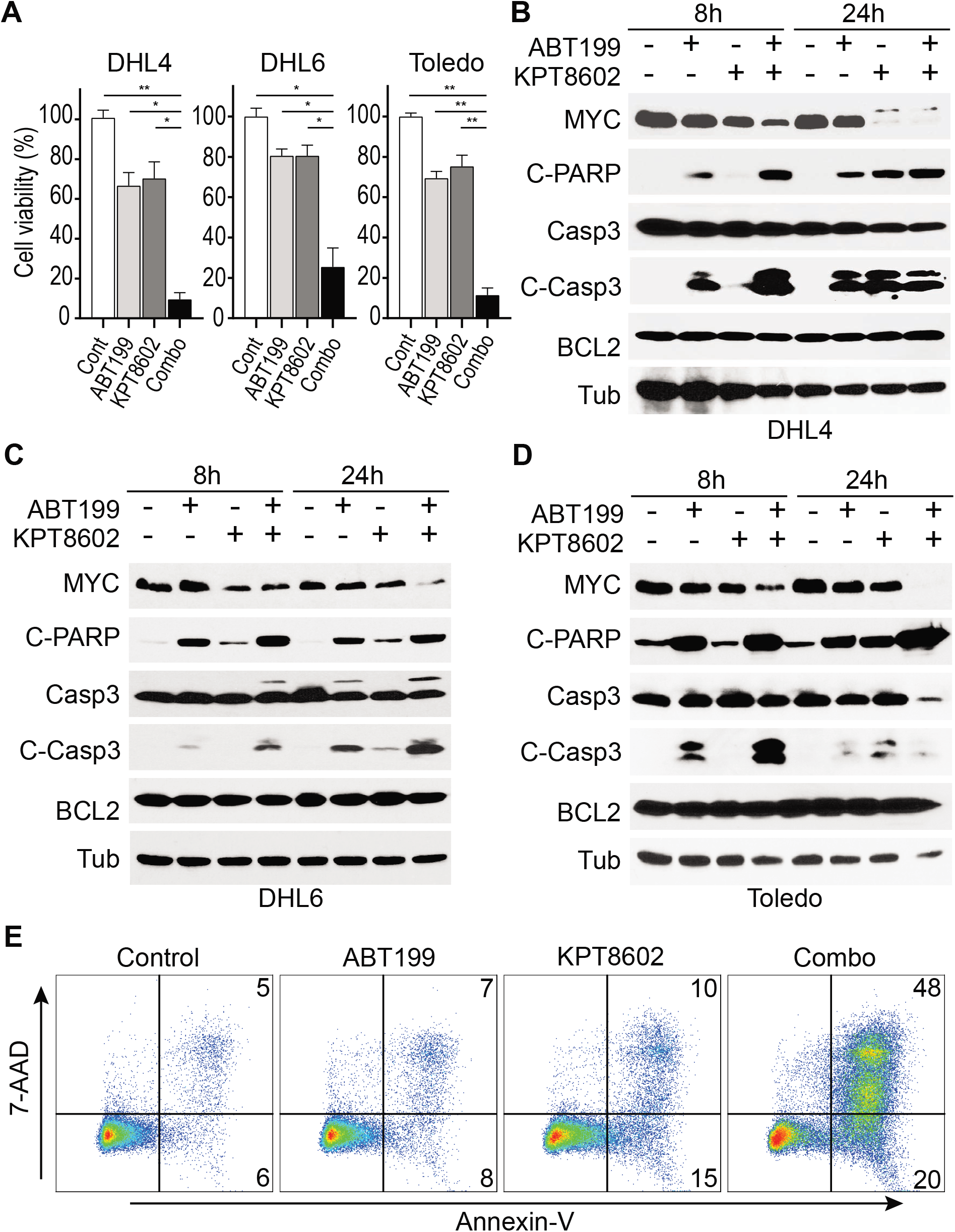
KPT8602 synergizes with ABT199 *in vitro* to kill DHL tumor cells. (A) Cell viability upon treatment with KPT8602 (100 nM) and ABT199 (40 nM for DHL4/DHL6, and 20 nM for Toledo) for 48 hours. (Two-tailed test. ** indicates P< 0.0001; * indicates P< 0.001.) (B-D) DHL cells were treated with 80 nM ABT199, 100 nM KPT8602 or both for 8 or 24 hours. Western blot analysis of protein lysates for MYC, cleaved PARP (C-PARP), Caspase3 (Casp3), cleaved Caspase3 (C-Casp3), BCL2, and β-Tubulin (Tub). (E) Flow cytometric analysis of apoptosis induction in DHL4 treated with 100 nM ABT199 and 50 nM KPT8602 for 48 hours. Cells were stained with Annexin-V and 7-AAD.

### Combined targeting of XPO1 and BCL2 blocks tumor progression and spread in vivo

Encouraged by our *in vitro* results, we initiated an *in vivo* study to test the combination therapy. Primary DHL patient-derived xenograft (PDX) tumor cells obtained from the Public Repository of Xenografts (PRoXe) were transplanted in NOD/SCID/IL2R gamma (NSG) mice [21]. These PDX tumor cells were labeled with firefly luciferase for bioluminescence imaging (BLI). Importantly, as part of the therapeutic regimen, a low dose of KPT8602 (7.5mg/kg, half of dose used in other studies) was used [13–15]. Additionally, at the start of the treatment, the tumor loads were at least 5 folds higher than recommended [21, 22]. Despite the low drug dose and higher than recommended tumor loads, we obtained a significant therapeutic response. The combined treatment completely blocked tumor progression and significantly extended survival, while single agents were only moderately effective (**Fig 4A-C**). Importantly, BLI signals from the skulls were reduced by 22 folds indicating the drug treatment effectively blocks tumor metastasis to brain (**Fig 4B, Supplementary Fig S5**). Tumor samples were also collected 5 days after drug treatment for immunohistochemical analysis of MYC expression, tumor cell proliferation, and apoptosis (Ki67 and cleaved Caspase 3). The combined drug treatment drastically abrogated MYC protein expression, decreased tumor cell proliferation and induced apoptosis (**Fig 5**). Thus, combined targeting of XPO1 and BCL2 is highly effective for the treatment of human DHL *in vivo*.

**Figure 4.**
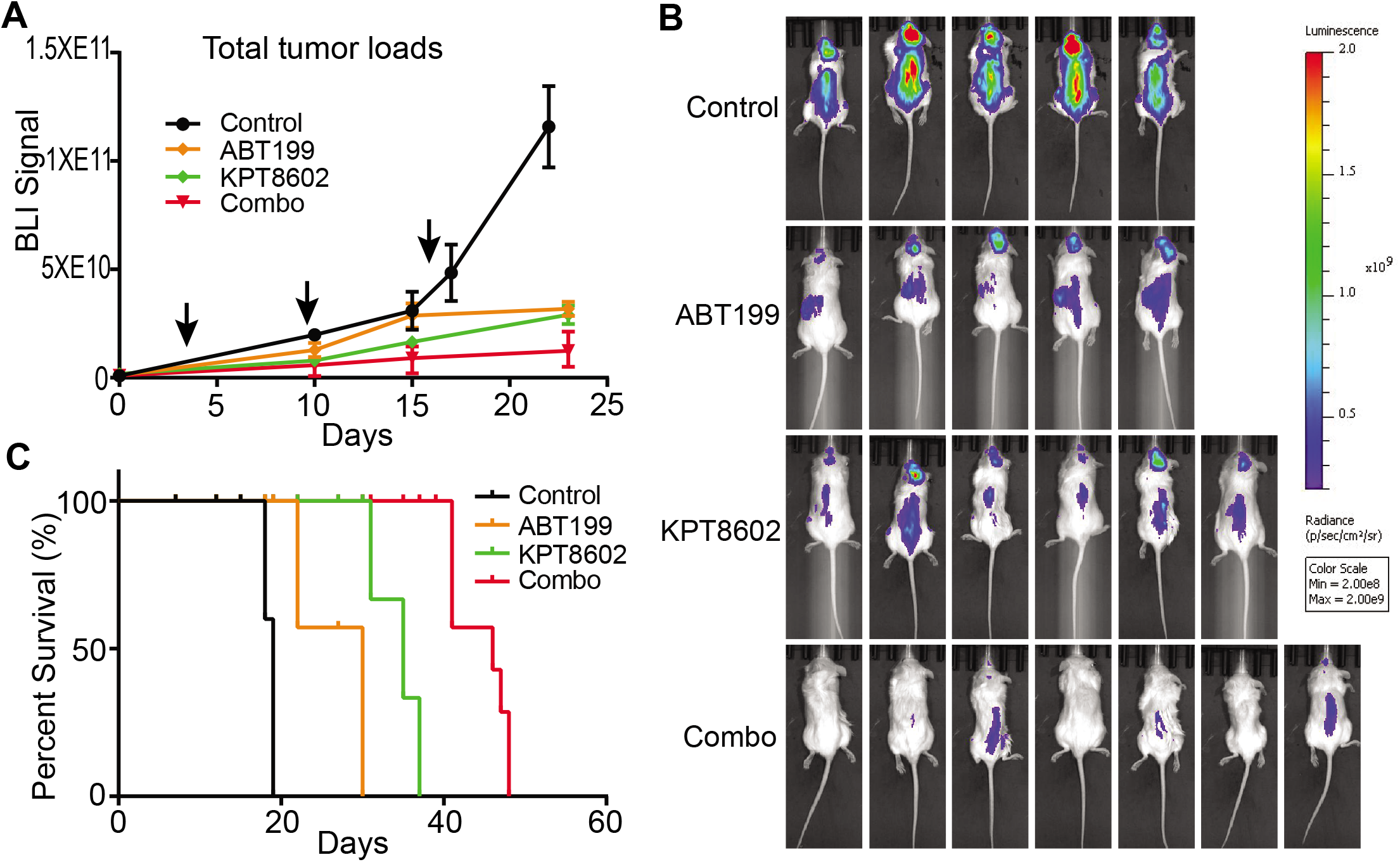
*In vivo* therapeutic effects of combining XPO1 and BCL2 inhibitors. (A) PDX tumor growth in NSG mice monitored by BLI. Arrows indicate three cycles of treatment (N=5, 5, 6, and 7 for control, ABT199, KPT8602, and combination groups, respectively). BLI data were presented as mean + standard error of mean. (B) Bioluminescent images of tumor-bearing mice following drug treatments. Control mice were imaged 15 days post-treatment. Drug-treated mice were imaged 21 days post-treatments. (C) Kaplan-Meier survival analysis of tumor-bearing mice (Log-rank Mantel-Cox test. P<0.0001).

**Figure 5.**
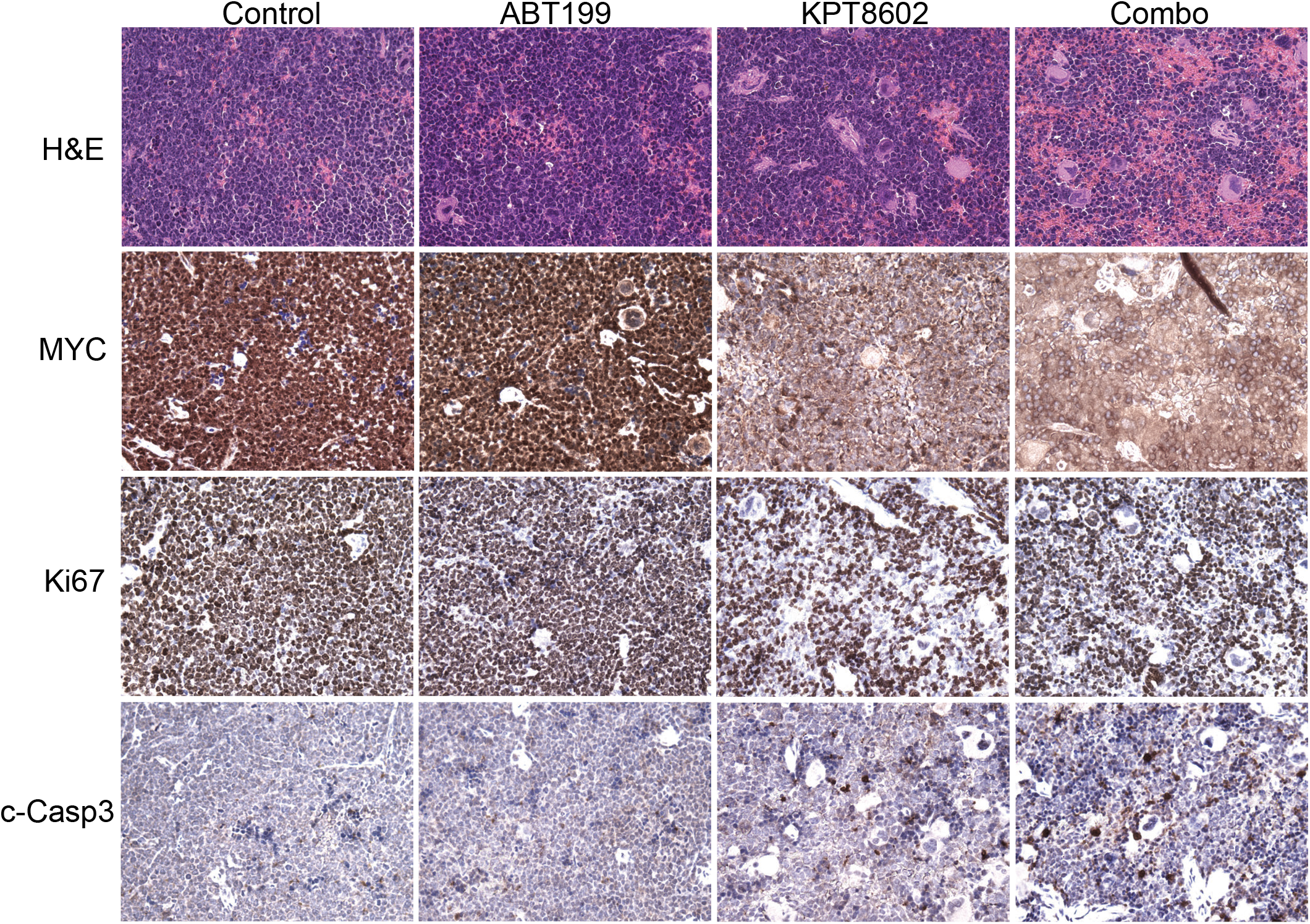
Immunohistochemical analysis of MYC expression, proliferation and survival of tumor cells. Spleen tumor samples were collected following a five-day treatment (one cycle). Ki67 and cleaved Caspase 3 (c-Casp3) staining were used to assess proliferation and survival of tumor cells.

## Discussion

XPO1 inhibition is a potential strategy to target MYC oncogene in DHL tumors as XPO1 inhibitors effectively downregulate MYC protein expression and reprogram the gene expression of MYC downstream targets. The mechanism by which XPO1 inhibition abolishes MYC protein expression remains unknown. We found that neither the expression nor the transport of MYC transcript can explain the downregulation of MYC expression by XPO1. Protein degradation as a contributing factor was also ruled out as proteasome inhibition did not rescue the downregulation of MYC by XPO1 inhibitors. These observations point to a translational regulation of MYC expression by XPO1. MYC mRNA has very long 5’ and 3’UTRs, containing regulatory motifs (such as IRES and ARE) important for translational initiation. We speculate that XPO1 may control nuclear cytoplasmic partitioning of key translation initiation factors essential for MYC protein expression. A comprehensive proteomic and genetic approach is underway to identify the *bona fide* translational regulator(s) in DHL tumor cell fractions.

In DHL tumors carrying genomic translocations of MYC and BCL2, and overexpressing the two driver oncogenes, MYC and BCL2 may be targeted simultaneously using XPO1 and BCL2 inhibitors. Accordingly, we find that concomitant suppression of XPO1 and BCL2 is effective for treatment of human DHL tumors. XPO1 and BCL2 inhibitors synergize *in vitro* to induce apoptosis in DHL tumor cells, and most importantly, block tumor progression and dissemination *in vivo*. Although our study has focused on the treatment of DHLs with genomic translocation of MYC and BCL2, the concept of simultaneously targeting multi-driver oncogenes, may be extended to other DHLs or triple-hit lymphomas (THLs) with genomic abnormalities in BCL6 oncogene. The inclusion of BCL6 inhibitors, such as 79-6 [23], in our combined therapeutic regimen could be tested in preclinical settings for the treatment of DHL/THLs.

Compared to other types of B-cell lymphomas, DHL tumors are more likely to spread to a patient’s central nervous system (CNS) [24]. Brain involvement in lymphoma patients generally confers a more abysmal prognosis with a median survival of 2-5 months [24]. Our *in vivo* study with human PDX reveals that combined targeting of XPO1 and BCL2 blocks brain metastasis of DHL tumors. Thus, it may be possible to prevent tumor spread using this type of combination targeted therapy. Our finding is in concordance with a recent case report, where the use of another XPO1 inhibitor, selinexor (KPT330), was shown to restrain the CNS relapse in one DLBCL patient [25]. The role of XPO1 in brain metastasis and the notion of using XPO1 inhibitors to suppress CNS involvement of lymphoma may warrant further preclinical examination.

## Supplementary Figure Legends

**Figure S1**

(A) Western blot analysis of MYC, MCL1, XPO1, and PARP/Caspase 3 cleavage. A panel of DHL cell lines was treated with 1 μM KPT8602 for 24 h. Part of the data was shown in **Fig 1**.

(B) Western blot analysis of MYC and PARP/Caspase 3 cleavage upon XPO1 inhibition in three DHL cell lines treated for 4, 8, 12, and 24 hours at drug concentration of 0.5 μM for DHL4/Toledo and 1 μM for DHL6.

(C) Western blot analysis of MYC and PARP/Caspase 3 cleavage upon treatment with different concentrations of XPO1 inhibitors for 24 hours in three DHL cell lines.

**Figure S2**

(A) Total mRNA and (B) nuclear to cytoplasmic ratios of BCL2, BCL6 and β-tubulin (internal control) in DHL6 treated with 1 μM KPT8602 for 4, 16 or 24 h. All mRNA levels were normalized to GAPDH.

(C) Quantification of nuclear and cytoplasmic Neat1 mRNA levels by realtime PCR.

(D) Analysis of nuclear and cytoplasmic GAPDH (cytoplasmic marker) and Lamin B1 (nuclear marker) by Western Blot.

(E) Representative DHL cells were treated with 1 μM KPT8602 and/or 10 nM Carfilzomib for 24 hours. β-actin was used as a loading control.

**Figure S3**

(A) IC50 values for ABT199 in a panel of DHL cell lines.

(B) IC50 values for KPT8602 in a panel of DHL cell lines.

(C) Cell viability in DHL cells treated with KPT8602 and ABT199 for 72 hours. The IC50 values for KPT8602 were calculated in the presence of different concentrations of co-administered ABT199.

(D) Cell morphology of DHL cells treated with KPT8602 (100 nM) and ABT199 (40 nM for DHL4/DHL6, and 20 nM for Toledo) for 48 hours.

**Figure S4**

Western blot analysis of MCL1, BCL-XL, and BIM proteins in DHL4 (A), DHL6 (B), and Toledo (C) cells. The drug treatment is the same as **Fig 3B-D**.

**Figure S5**

Quantification of BLI signals from the crania of the tumor bearing animals following drug treatment. BLI signal data were presented as mean + standard error of mean. N=5, 5, 6, and 7 for control, ABT199, KPT8602, and combination groups, respectively. The difference in BLI signals between control and single or combination treated mice was significant. Two-tailed t test. * Control vs ABT199, P=0.02; ** Control vs KPT8602, P=0.01; *** Control vs Combination, P=0.0008..

## Abbreviations

DHL: Double-hit lymphoma
NHL: Non-Hodgkin’s Lymphoma
PRoXe: Public Repository of Xenografts
(PDX): Patient-derived xenograft
BLI: Bioluminescence imaging
NSG: NOD/SCID/IL2R gamma
CNS: Central nervous system

## Declarations

### Ethics approval and consent to participate

Animal studies and experimental protocols were approved by Institutional Animal Care and Use Committee at Houston Methodist Research Institute. All experimental methods were performed in accordance with the relevant national and institutional guidelines and regulations. The DHL PDX sample was obtained from the Public Repository of Xenografts (PRoXe), which collects PDX samples from patients with informed consent.

### Consent for publication

Not applicable

### Availability of data and material

Not applicable

### Authors’ contributions

YLiu performed the *in vitro* experiments. NAzizian performed the *in vivo* therapeutic experiments. YDou and Yliu performed the flow cytometric analysis. LVPham provided scientific inputs and edited the manuscript. YLi conceived the study and wrote the manuscript. All authors read and approved the final manuscript.

## Acknowledgements

Not applicable

