## Supplementary figures and images for "Simultaneous Targeting of XPO1 and BCL2 as an Effective Treatment Strategy for Double-Hit Lymphoma"

### Supplementary Fig S1-S5

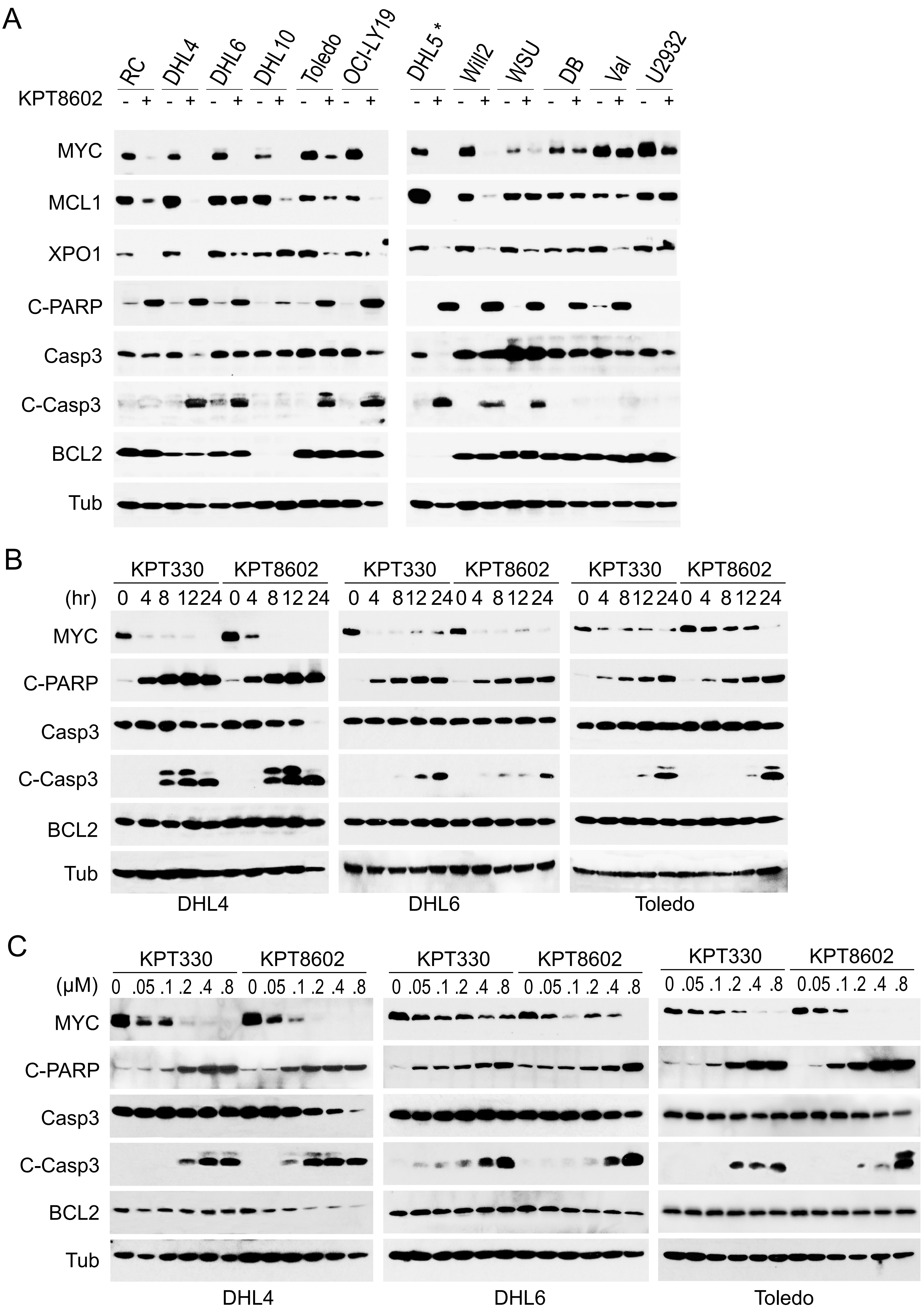

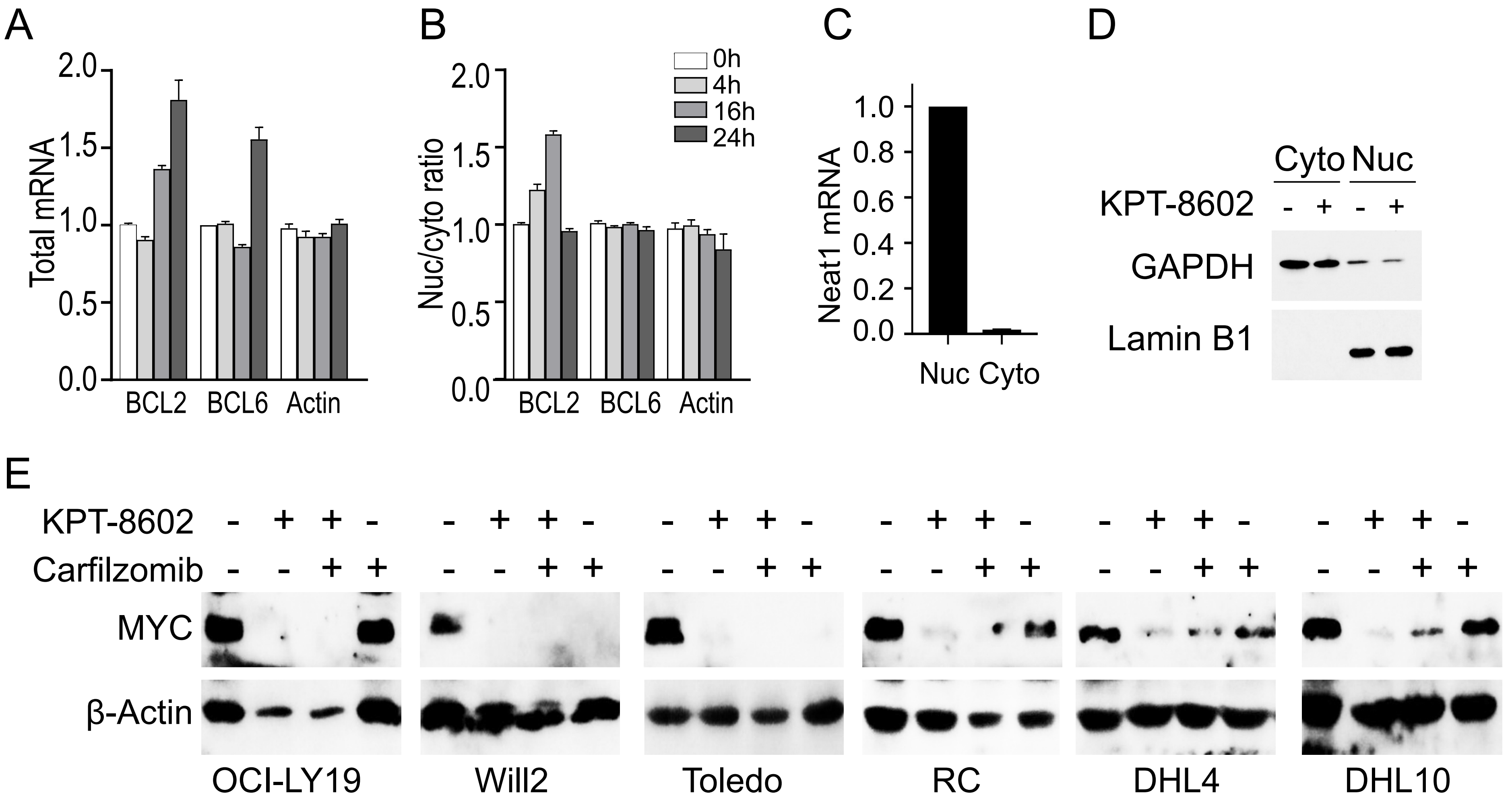

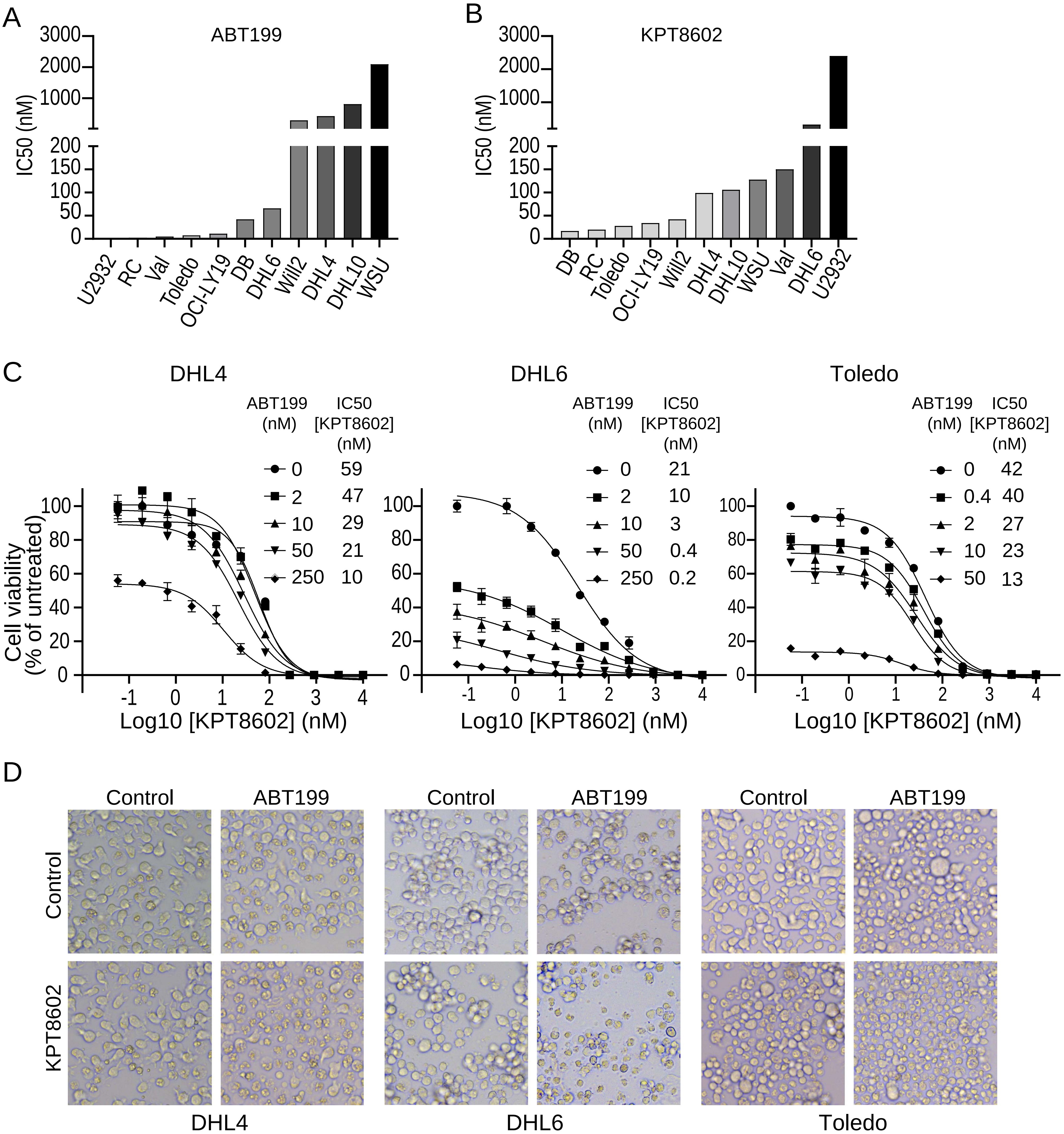

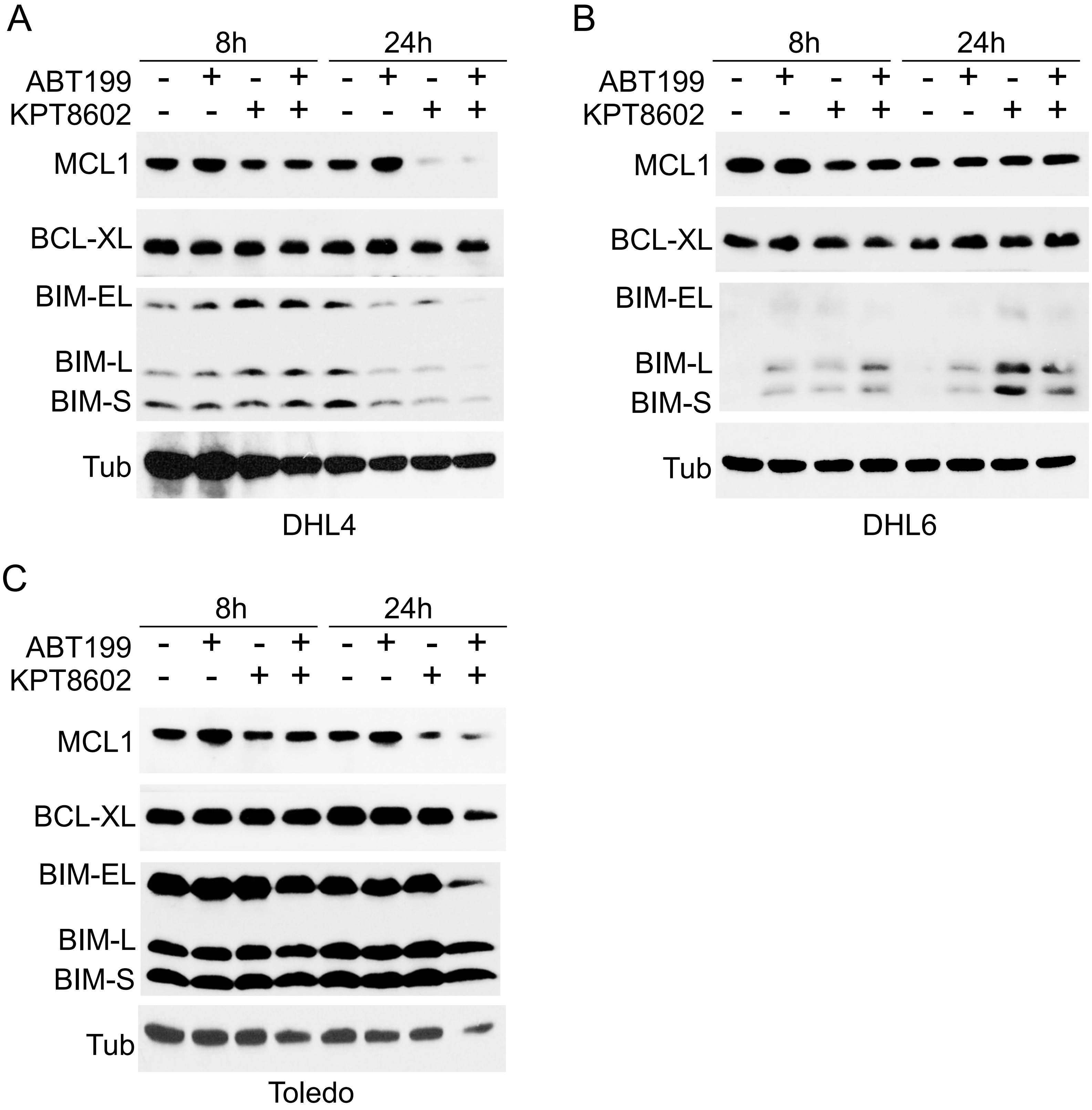

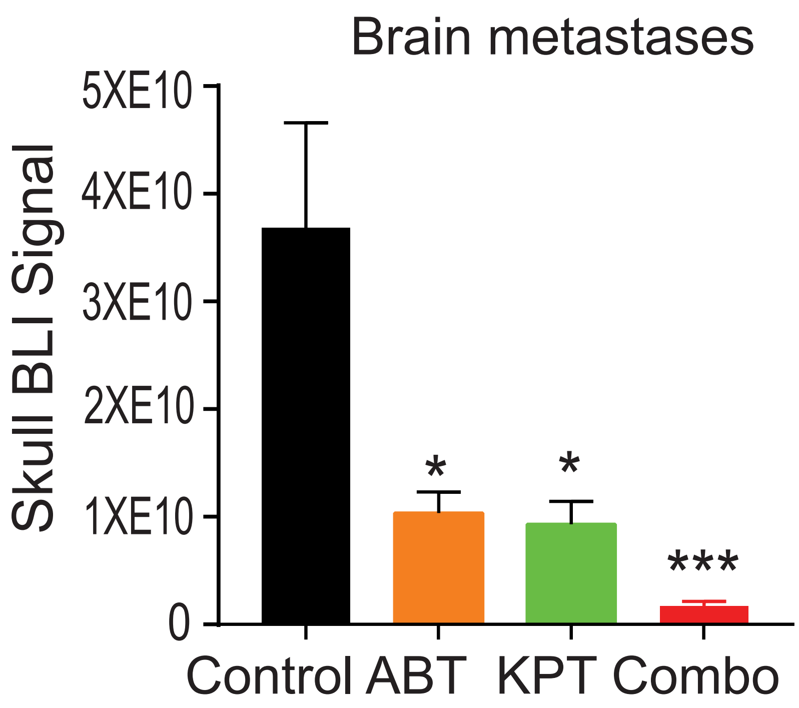
